# Does *Borrelia afzelii* outer surface protein E coevolve with complement factor H of its rodent host? Insights from GxG and spatial associations

**DOI:** 10.64898/2026.07.10.737716

**Authors:** Joanna Różańska-Wróbel, Karolina Przesmycka, Józefina Wasilewska, Maciej Grzybek, Rocco F. Notarnicola, Anna Bajer, Dorota Dwużnik-Szarek, Mohammed Alsarraf, Jolanta Behnke-Borowczyk, Jerzy M. Behnke, Jacek Radwan

## Abstract

**Background:** Lyme borreliosis is a common tick-borne disease in Europe caused by spirochetes of the *Borrelia burgdorferi* sensu lato complex, including *Borrelia afzelii*, which is maintained in nature through interactions with rodent reservoir hosts. These spirochetes have evolved several surface proteins to manipulate rodent host immunity, some of which remain polymorphic in *Borrelia* populations. Among these proteins, OspE, which binds the host complement-regulating factor CFH to evade destruction by complement, is one of the most variable. Yet, what evolutionary forces maintain this polymorphism is not well understood. Motivated by a recent discovery of CFH polymorphism in the bank vole (*Clethrionomys glareolus*), the main reservoir host of *B. afzelii*, we hypothesized that the polymorphism is maintained by host–parasite coevolution involving specific associations between host and parasite genetic variants.

**Methods:** We analyzed associations between bank vole CFH alleles and *B. afzelii* OspE variants across three datasets sampled in Poland. Selection acting on OspE was evaluated using omegaMap. Host-pathogen genotype associations were tested using partial redundancy analysis (RDA), and co-structure was assessed using co-correspondence analysis (CoCA).

**Results:** We found that OspE evolves under positive selection, however, we found no evidence for an association between OspE and host CFH variants at the individual level based on RDA or at the population level based on CoCA.

**Conclusions:** Despite evidence of positive selection acting on OspE, we found no support for specific genetic matching between *B. afzelii* and its bank vole host at the CFH–OspE interface. These results suggest that the evolution of CFH and OspE may be shaped by broader selective pressures, potentially including interactions with multiple host species.

## Background

Parasites evolve various ways of evading host immune defenses, including antigenic variability, hiding from immune surveillance, complement inhibition, and the direct subversion or destruction of host immune cells [1,2]. This, in turn, selects for host counter-adaptations, leading to either a series of recurrent selective sweeps that result in the fixation of advantageous alleles (the “arms race” model) or the maintenance of high genetic diversity via negative frequency-dependent selection (the “balancing selection” model) [3]. Both coevolutionary scenarios assume that specific variants of parasite infectivity factors differentially infect hosts carrying alternative immune alleles, leading to transient or stable polymorphisms, respectively. Investigating whether such specific genetic interactions occur provides an empirical test of these coevolutionary models. Furthermore, understanding how parasite polymorphisms evolve has critical consequences for epidemiology and vaccine development [4–6].

*Borrelia burgdorferi* sensu lato (Bbsl henceforth), an agent of Lyme disease, has evolved several outer surface proteins to overcome host immune defenses [7]. These include the continuous antigenic variation of the VlsE lipoprotein to escape adaptive immune recognition [8], as well as the expression of OspC, which facilitates early dissemination and evasion of innate phagocytosis [9]. Furthermore, *Borrelia* produces a suite of complement regulator-acquiring surface proteins (CRASPs) that hijack host complement regulators, such as complement factor H (CFH), to inhibit complement-mediated lysis [10]. OspE is among the most variable of these proteins, and yet the mechanisms maintaining this polymorphism are not well understood.

The OspE lipoprotein, together with OspF and Elp proteins, belongs to a family known as Erp (OspEF-related) proteins, which are encoded on the circular cp32 plasmids of *Borrelia* [11]. Each single bacterium may carry multiple copies of cp32 plasmids and express different OspE proteins simultaneously [12], and *Borrelia* strains vary in OspE composition [13–15]. In contrast, another surface protein, encoded by the single-copy ospC gene, is commonly used as a marker of *Borrelia* strain identity [16], allowing infecting strains to be inferred from OspC variants and enabling comparison of associated OspE variant repertoires. OspE protein-coding regions are highly diverse, with the diversity promoted by intergenic recombination events with closely related as well as distantly related Erps [13]. OspE proteins are exposed on the surface of *Borrelia*, and their expression is upregulated during tick feeding and maintained throughout mammalian infection [17]. Their main function appears to be avoidance of destruction by the host’s complement system. Specifically, OspE binds to the CCP20 domain of host CFH, a key regulatory protein of the alternative complement pathway [18,19]. Unlike the classical and lectin pathways that are activated by signals of infection, the alternative pathway does not require specific pathogen-binding proteins but is instead constitutively active at a low level and is amplified on non-self surfaces. The regulatory protein CFH inactivates the alternative pathway on the surface of host cells by preferentially binding to vertebrate cells via polyanions, such as sialic acid, present on those cells but absent from pathogen surfaces. Binding of CFH inhibits the C3 convertase formation and activity of the alternative pathway, thereby preventing complement-mediated damage of host cells [20,21]. The binding of CFH to the surface of *Borrelia* therefore protects the pathogen from complement-mediated destruction and thus enables its survival in host serum [22].

Interestingly, Notarnicola et al. [23] found that CFH is polymorphic in one of the key rodent reservoirs of Bbsl in Europe, the bank vole (*Clethrionomys glareolus*). Furthermore, analysis of CFH sequences across cricetid and murid rodents has indicated that CFH evolved under positive selection, with the CCP20 domain that binds to Bbsl showing the highest number of polymorphic codons under selection [23]. Likewise, analyses of several *Borrelia* species have revealed signatures of positive selection at codons interacting with CFH [24], suggesting that CFH and OspE may coevolve tightly. Here, we hypothesized that within-species polymorphism of CFH may be driven by its coevolution with OspE of the most common Bbsl species in Europe, *B. afzelii*. Specifically, we expected that *B. afzelii* strains carrying specific OspE variants would be associated with infection of hosts carrying specific CFH. Furthermore, we hypothesized that such associations could drive the CFH genetic population structuring reported for bank vole populations; Notarnicola et al., 2026 found that the genetic structure of CFH across 13 populations of bank voles in Poland was stronger than the genomic average, implying that local adaptation reduces the flow of CFH variants between populations. These local pressures could include adaptation to local *B. afzelii* genotypes, in which case host population structure should mirror the distribution of *Borrelia* OspE variants, and could be shaped by *Borrelia* interactions with multiple host species [25,26]. However, little is known about OspE diversity and the signatures of molecular evolution within *B. afzelii*. Furthermore, to our knowledge, population structuring of OspE has not been investigated in any Bbsl species. Published studies of population structure based on other markers [27,28] may not reflect the structure of OspE due to extensive horizontal transfer reported for cp32 plasmids [29,30]. Therefore, to understand how host-parasite coevolution shapes polymorphisms in CFH and OspE, known to interact at the molecular level [18,31,32], we investigated (i) whether OspE in *B. afzelii* evolves under positive selection at sites known to interact with CFH, and (ii) whether specific variants of the bank vole CFH gene family are associated with infection by *B. afzelii* strains carrying different OspE gene variants in populations polymorphic for CFH. Furthermore, we investigated if OspE shows significant genetic structuring and if this structuring results in a spatial co-structure between OspE and CFH across populations in Poland.

## Methods

### Bank vole samples and determination of their infection status

This research used three sets of samples. The first two were used to investigate the association between infection with *B. afzelii* strains carrying particular OspE variants and bank vole CFH genotypes within populations, and the third to investigate OspE population genetic structure and its co-structure with CFH. The first two sets were analyzed separately because they corresponded to different sampling periods and populations, and the second set included additional CFH variants not captured in the first set.

The first set consisted of the same bank vole samples as in our previous study of vole MHC and *Borrelia* OspC genes [28]. In brief, bank voles were sampled from 2002 to 2018 at 4-year intervals in Northeastern Poland at three study sites spaced approximately 10 km apart: Urwitałt, Tałty and Pilchy (Table S1). Trapping and sampling were performed under the approval of the First Warsaw Local Ethics Committee for Animal Experimentation. Sex and age class were assigned as described in Różańska-Wróbel et al. [28], tail samples were collected, stored at −20°C, and total DNA was extracted using the MagJET Genomic DNA Purification Kit (Thermo Fisher Scientific, Waltham, MA, USA). Infection status with *B. afzelii* was established as described in Różańska-Wróbel et al. [28].

The second set consisted of bank vole ear biopsy samples collected between August and October 2023-2025 from three populations selected for their high CFH diversity (Brok, Grobka, Długosiodło; Table S2). These populations were also included in the third dataset. The third set, used for OspE genetic structure and co-structure analysis, consisted of vole ear biopsy samples collected from 12 populations across Poland (Table S3) between August and October 2020-2023.

Sampling for the second and third sets was conducted using live trapping, under the approval of the Local Ethics Committee for Animal Experimentation in Poznań, decision no. 35/2021. Following sampling, all voles were weighed and released at the site of capture. Ear samples were stored in ethanol at −20°C, and total DNA was extracted using the NucleoMag 96 Tissue Kit (Macherey-Nagel, Duren, Germany). Bank vole sex was determined using PCR as described in Różańska-Wróbel et al. [33]. *B. afzelii* infection was detected using primers targeting the rrs-rrl (16S-23S) ribosomal intergenetic spacer [34]. Nested PCRs were run with the Type-it Microsatellite PCR Kit (Qiagen, Hilden, Germany) for 30 cycles at an annealing temperature of 52°C, with 1 μL of template and 0.4 μM of each primer.

### CFH and OspE genotyping

Bank vole CFH 227–245 bp fragment, covering most of the CCP20 domain, was amplified using primers CCP20-F: 5’-CATGTGTAATATCAGAAGAGAYCA-3’ and CCP20-R: 5’- CATGAATATTTCTMAACAAAGGTTCAT-3’, described by Notarnicola et al. [23]. The primers also amplify some factor H-related (FH-R) fragments, which were distinguished from CFH as described below. The primers were barcoded, and the ‘N’ (0-3 nucleotide) spacers were added to increase library diversity during sequencing. The barcodes used to identify unique samples have been described previously by Kozich et al. [35]. PCRs were performed using the Type-it Microsatellite PCR Kit according to the manufacturer’s protocol. Each reaction mixture (total volume of 25 µL) contained 1 µL of template DNA, 1 µL of CCP20-F primer (final concentration: 0.4 µM), 1 µL of CCP20-R primer (final concentration: 0.4 µM), 12.5 µL of 2x Type-it Multiplex PCR Master Mix, and nuclease-free water up to 25 µL. Annealing temperature was set to 58°C and the PCR was run for 28 cycles.

*B. afzelii* OspE gene fragments were amplified from the infected bank voles using barcoded primes Baf_ospE-F: 5’-GCAATGACTRAAGGYGGATCA-3’ and Baf_ospE-R: 5’-TCTTCTARTGRTATTGCAT-3’, designed based on alignment of available *B. afzelii* sequences, co-aligned with *B. burgdorferi* sensu stricto (s.s.) to identify conserved regions. The length of the PCR product was 200–272 bp and included a region known to interact with CFH [18,24]. PCRs were performed using the Type-it Microsatellite PCR Kit. Each reaction mixture (total volume of 25 µL) contained 1 µL of template DNA, 1 µL of Baf_ospE_F primer (final concentration: 0.4 µM), 1 µL of Baf_ospE_R primer (final concentration: 0.4 µM), 12.5 µL of 2x Type-it Multiplex PCR Master Mix, and nuclease-free water up to 25 µL. Annealing temperature was set to 56°C and the PCR was run for 32 cycles.

Sequencing libraries were prepared using the NEBNext Ultra II DNA Library Prep Kit for Illumina and NEBNext Multiplex Oligos for Illumina (New England Biolabs, Ipswich, MA, USA). The CFH and OspE amplicons were sequenced on the Illumina MiSeq platform using the MiSeq Reagent Micro Kit v2 (paired-end, 2 × 150) in separate runs. Obtained paired-end reads were merged with the AmpliMERGE tool and genotyped with the AmpliSAS software using the default Illumina parameters for de-multiplexing, clustering and filtering of unique variants [36].

CFH and FH-R sequences were then mapped to a reference database of expressed CFH obtained from cDNA sequencing, as described in Notarnicola et al. [23]. In subsequent analyses, we focused on sequences matching expressed CFHs, as the function of FH-R variants is not well understood [37]. Nevertheless, we also carried out parallel exploratory analyses that included both CFH and FH-R variants.

### Statistical analyses

To detect selection at individual sites of the OspE gene variants found in the studied populations, we used omegaMap, v. 0.5 [38], which estimates the ratio of nonsynonymous to synonymous substitutions (ω) using a Bayesian framework while accounting for recombination (ρ). We analyzed a 132 bp-long OspE fragment encoding 44 amino acids (positions 106-149), which was found to contain positively selected sites involved in protein interactions at binding sites [24]. Sequences were aligned using MUSCLE in AliView, v. 1.30 [39]. Analyses were performed on the aligned sequences using an independent ω model and a variable ρ model with a block size of 20. Codon frequencies were assumed equal (i.e. 1/61). Priors were set as improper inverse for μ, κ, and ϕ, and an inverse prior for ω and ρ. Markov Chain Monte Carlo (MCMC) was run for 2,000,000 iterations with thinning every 100 iterations, using 10 orderings and a burn-in of 10,000. Three independent runs of omegaMap were performed to ensure convergence and were visualized in R, v. 4.1.2 [40], using Rstudio, v. 2024.4.2.764 [41] by plotting the ω point estimates along with the lower and upper bounds of the 95% highest posterior density (HPD) intervals against codon position. The results of these runs were then combined to generate the posterior distributions. Codons were considered to be under positive selection if the posterior probability that ω > 1 was greater than 0.95.

Potential associations between bank vole CFH variants and *B. afzelii* OspE variants were examined with a multivariate statistical method, the redundancy analysis (RDA). We performed partial RDA with the rda function from the vegan R package, v. 2.6-4 [42]. CFH alleles were used as explanatory variables; sampling year, site, host age (dataset 1) or body mass (dataset 2) and sex were included as covariates; and the presence or absence of OspE variants were used as response variables. To reduce sparsity and improve the stability of multivariate models, we excluded low-frequency variants prior to analysis. Because CFH variants were used as explanatory variables, we applied a relative frequency threshold, retaining only those present in >10 % of bank voles to prevent overparameterization and ensure stable coefficient estimates. OspE variants (response variables) were filtered using an absolute threshold, retaining only variants detected in at least 10 bank voles. This approach removes uninformative rare occurrences while ensuring sufficient observations per variant for stable model fitting and preserving meaningful variation for ordination. The significance of the RDA models was assessed with permutation tests conducted with the anova.cca function from the vegan package. The results of permutation tests were averaged over 50 runs of anova.cca.

To examine the genetic structure of the OspE gene across populations, a haplotype network was constructed using the haploNet function from the pegas R package, v. 1.2 [43]. Haplotype frequencies across populations were calculated using the haploFreq function. The network was visualized with node sizes proportional to haplotype frequencies and pie charts representing the relative contribution of each population.

To predict the composition of bank vole CFH genes from *B. afzelii* OspE composition, predictive co-correspondence analysis (CoCA) was performed using the cocorresp R package, v. 0.4-6 [44]. Log-transformed OspE variant abundances in each population were used as the predictor matrix, and log-transformed CFH variant abundances in each population as the response matrix. Predictive accuracy was evaluated with leave-one-out cross-validation using the crossval function. A positive cross-validatory fit indicates that the model predicts the response data better than chance. The significance of each CoCA axis was evaluated with permutation tests using the permutest function from the vegan package, averaged over 50 runs.

## Results

In total, we identified 44 OspE variants: 36 variants in our large sample from the northeastern subpopulations (Uwritałt, Tałty and Pilchy), including eight rare variants not found in the second and third datasets (each found in no more than three individuals), and eight additional sequences found exclusively in the remaining sites. The phylogenetic tree of the detected OspE variants, along with *B. burgdorferi* Erp sequences downloaded from the GenBank database, is shown in Figure 1. All *B. afzelii* OspE sequences grouped with *B. burgdorferi* Erps A, C and P.

**Fig. 1.**
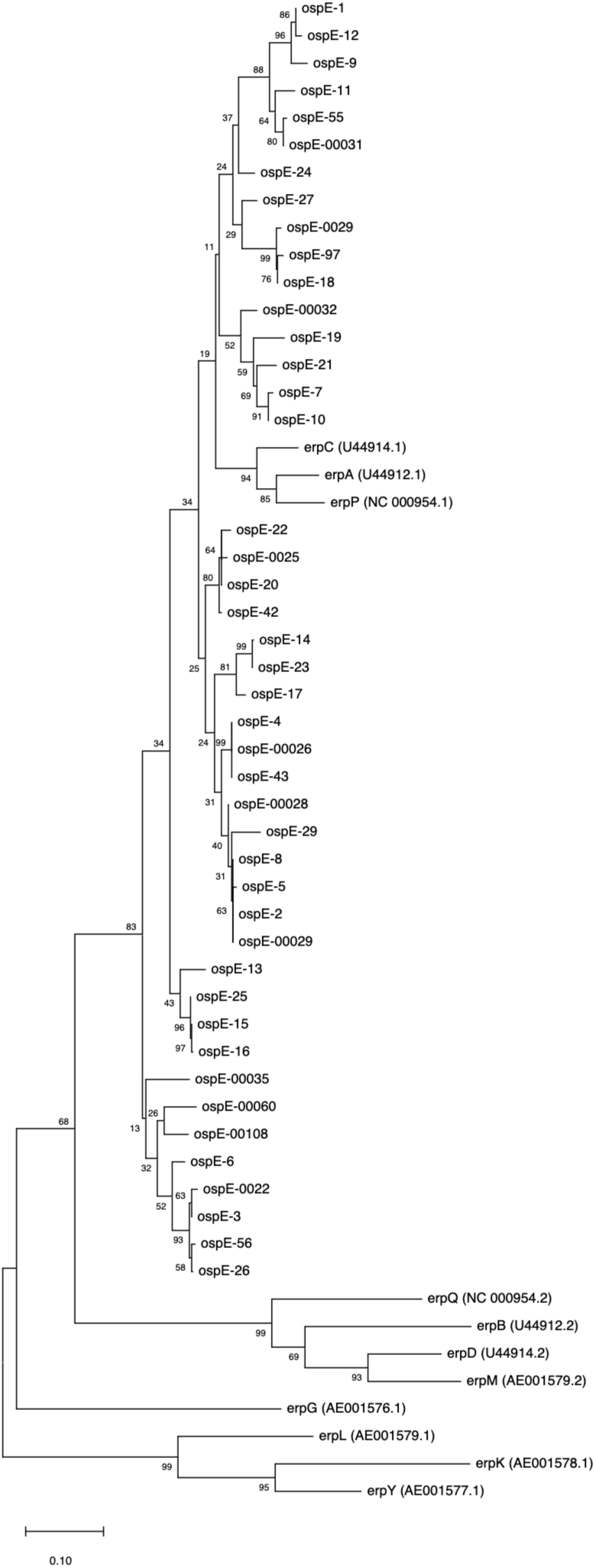
Neighbor-Joining tree of *B. afzelii* OspE variants detected in all analyzed bank vole samples (names starting with “ospE”), along with *B. burgdorferi* Erp sequences downloaded from GenBank (names starting with “erp”; accession numbers shown in parentheses). Evolutionary distances were computed using the Maximum Composite Likelihood method. Bootstrap support values (based on 500 replicates) are shown at the nodes.

Among infected individuals, we identified 60 single-strain infections inferred from the presence of only one OspC variant [28]. These strains harbored one to six OspE variants. A histogram of OspE variant counts in single-strain infections is shown in Figure S1, and a haplotype combination showing OspE variant co-occurrences within these strains is shown in Table S4. While strains carrying the same OspC sometimes shared OspEs that were absent or very rare among strains carrying different OspCs (e.g. OspE2-OspC10b, OspE6-OspC7b, OspE20-OspC3a), most OspC strains had multiple OspE combinations, and strains carrying different OspC types often shared the same OspEs (Table S4).

We found evidence of recombination and positive selection acting on the identified OspE variants (Table 1). The three independent runs of omegaMap showed good convergence (Figure S2). After combining the runs, the posterior distributions identified eight codon sites under positive selection with Pr(ω > 1) ≥ 0.95 at codons 116, 119, 120, 121, 122, 138, 139, and 145 (Table 1; codon positions refer to amino acid residues in the full-length OspE protein sequence).

**Table 1.** Site-specific estimates of ω (dN/dS) and ρ, 95% highest posterior density (HPD) intervals, and posterior probabilities of positive selection for the analyzed OspE fragment inferred using omegaMap. Codons with Pr(ω > 1) ≥ 0.95 are shown in bold. Site numbers refer to amino acid residues in the full-length OspE protein sequence.

| Site | $\omega$ lower<br>95% HPD | $\omega$ point<br>estimate | $\omega$ higher<br>95% HPD | Posterior<br>prob. of<br>positive<br>selection | $\rho$ lower<br>95% HPD | $\rho$ point<br>estimate | $\rho$ higher<br>95% HPD |
| --- | --- | --- | --- | --- | --- | --- | --- |
| 106 | 0.010 | 0.129 | 1.981 | 0.123 | 0.010 | 0.522 | 2.249 |
| 107 | 0.010 | 0.150 | 2.614 | 0.156 | 0.013 | 0.504 | 1.647 |
| 108 | 0.117 | 1.620 | 19.179 | 0.683 | 0.023 | 0.487 | 1.631 |
| 109 | 0.010 | 0.143 | 2.556 | 0.150 | 0.029 | 0.500 | 1.651 |
| 110 | 0.010 | 0.159 | 3.342 | 0.164 | 0.038 | 0.519 | 1.651 |
| 111 | 0.010 | 0.189 | 3.375 | 0.201 | 0.053 | 0.545 | 1.637 |
| 112 | 0.010 | 0.154 | 2.572 | 0.156 | 0.081 | 0.574 | 1.635 |
| 113 | 0.010 | 0.159 | 3.098 | 0.171 | 0.115 | 0.599 | 1.658 |
| 114 | 0.098 | 1.520 | 17.756 | 0.665 | 0.138 | 0.613 | 1.641 |
| 115 | 0.688 | 4.045 | 26.082 | 0.927 | 0.164 | 0.627 | 1.643 |
| **116** | **1.983** | **9.841** | **51.640** | **0.997** | **0.293** | **0.678** | **1.631** |
| 117 | 0.010 | 0.128 | 2.428 | 0.138 | 0.348 | 0.712 | 1.507 |
| 118 | 0.010 | 0.161 | 3.417 | 0.179 | 0.373 | 0.740 | 1.421 |
| **119** | **2.148** | **11.086** | **58.025** | **0.997** | **0.390** | **0.750** | **1.426** |
| **120** | **6.654** | **23.035** | **85.558** | **1.000** | **0.390** | **0.751** | **1.407** |
| **121** | **6.807** | **23.784** | **90.186** | **1.000** | **0.394** | **0.753** | **1.408** |
| **122** | **1.991** | **11.308** | **63.984** | **0.995** | **0.393** | **0.764** | **1.407** |
| 123 | 0.010 | 0.154 | 2.942 | 0.163 | 0.397 | 0.762 | 1.406 |
| 124 | 0.084 | 1.597 | 15.837 | 0.683 | 0.401 | 0.750 | 1.407 |
| 125 | 0.594 | 4.656 | 28.022 | 0.933 | 0.401 | 0.748 | 1.405 |
| 126 | 0.010 | 0.133 | 2.576 | 0.138 | 0.390 | 0.745 | 1.368 |
| 127 | 0.010 | 0.162 | 2.833 | 0.175 | 0.390 | 0.743 | 1.370 |
| 128 | 0.010 | 0.167 | 3.031 | 0.183 | 0.389 | 0.741 | 1.370 |
| 129 | 0.010 | 0.132 | 2.230 | 0.127 | 0.389 | 0.740 | 1.372 |
| 130 | 0.138 | 1.922 | 23.166 | 0.717 | 0.388 | 0.739 | 1.371 |
| 131 | 0.010 | 0.139 | 2.936 | 0.156 | 0.384 | 0.735 | 1.370 |
| 132 | 0.010 | 0.129 | 2.148 | 0.124 | 0.379 | 0.730 | 1.370 |
| 133 | 0.081 | 1.280 | 13.742 | 0.625 | 0.372 | 0.718 | 1.392 |
| 134 | 0.010 | 0.130 | 2.173 | 0.129 | 0.363 | 0.707 | 1.424 |
| 135 | 0.228 | 3.140 | 27.894 | 0.840 | 0.334 | 0.689 | 1.424 |
| 136 | 0.010 | 0.141 | 2.562 | 0.146 | 0.321 | 0.676 | 1.483 |
| 137 | 0.010 | 0.126 | 2.385 | 0.132 | 0.295 | 0.670 | 1.479 |
| **138** | **1.508** | **7.912** | **41.580** | **0.987** | **0.291** | **0.669** | **1.554** |
| **139** | **2.878** | **13.045** | **53.944** | **1.000** | **0.280** | **0.691** | **1.636** |
| 140 | 0.010 | 0.141 | 2.453 | 0.138 | 0.260 | 0.722 | 1.821 |
| 141 | 0.329 | 3.721 | 32.624 | 0.868 | 0.257 | 0.768 | 2.637 |
| 142 | 0.010 | 0.169 | 3.000 | 0.177 | 0.243 | 0.753 | 4.363 |
| 143 | 0.010 | 0.168 | 2.887 | 0.170 | 0.072 | 0.676 | 5.893 |
| 144 | 0.010 | 0.125 | 2.280 | 0.128 | 0.014 | 0.540 | 1.763 |
| **145** | **1.050** | **8.056** | **49.424** | **0.976** | **0.015** | **0.386** | **1.431** |
| 146 | 0.010 | 0.164 | 3.089 | 0.187 | 0.014 | 0.336 | 1.344 |
| 147 | 0.010 | 0.130 | 2.428 | 0.136 | 0.014 | 0.311 | 1.342 |
| 148 | 0.010 | 0.139 | 2.282 | 0.145 | 0.013 | 0.299 | 1.327 |
| 149 | 0.099 | 1.743 | 21.896 | 0.703 |  |  |  |

For the first dataset (Urwitałt, Tałty, and Pilchy populations), we obtained complete data (host age, sex, CFH and OspE variants) for 1146 bank voles, of which 158 (13.79%) were infected with *B. afzelii* (Table S1). Among the 36 OspE variants detected, 22 were found in at least 10 bank voles. Among the 51 CFH/FH-R variants detected, five corresponded to the CFH alleles previously described by Notarnicola et al. [23], and the remaining 46 were classified as FH-R. Two CFH variants occurred in more than 10% of bank voles, and together with the 22 OspE variants present in at least 10 individuals, these variants were used in partial RDA analyses. We examined associations between bank vole and *B. afzelii* genotypes in both the full dataset and the subset of infected voles. However, we found no significant associations in either case (full dataset: adjusted R^2^ = 1.27e-4, P = 0.351; infected subset: adjusted R^2^ = 0.003, P = 0.205; Table 2). Additionally, we performed RDA analyses using jointly 12 CFH and FH-R variants occurring in more than 10% of voles as explanatory variables, but again found no significant associations (full dataset: adjusted R^2^ = 1.81e-4, P = 0.424; infected subset: adjusted R^2^ = -0.013, P = 0.897; Table S5).

**Table 2.**
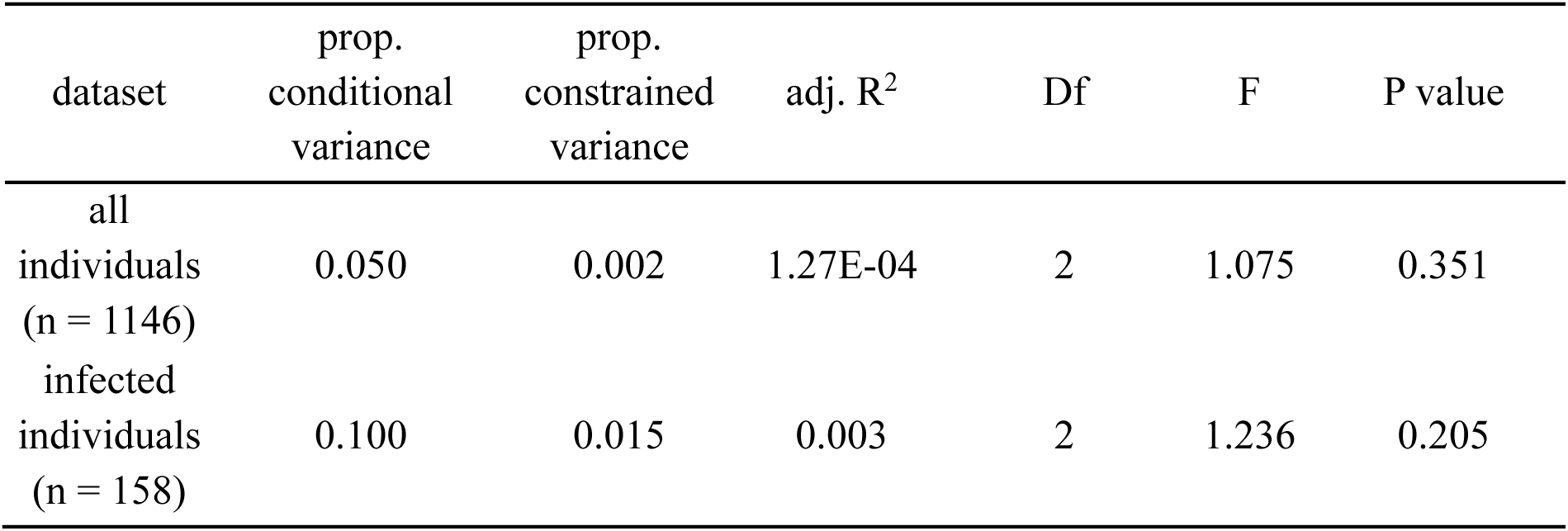
Results of partial redundancy analysis (RDA) performed on the first set of bank vole samples collected at three sites: Urwitałt, Tałty, and Pilchy. Bank vole CFH alleles were used as explanatory variables; sampling year, site, host age and sex were included as covariates; and the presence or absence of *B. afzelii* OspE variants was used as response variables.

For the second dataset (Brok, Grobka, and Długosiodło populations), we obtained complete data (host body mass, sex, CFH and OspE variants) for 411 bank voles, of which 41 (9.98%) were infected with *B. afzelii* (Table S2). In this dataset, 27 OspE variants were detected, with six present in at least 10 bank voles. A total of 48 CFH/FH-R variants were identified, including eight CFH alleles, five of which occurred in more than 10% of voles.

Here, too, no significant associations were observed between CFH and OspE in the full dataset (adjusted R^2^ = -4.09e-4, P = 0.492; Table 3). RDA using 17 CFH/FH-R variants present in > 10% of voles also revealed no significant associations (adjusted R^2^ = 7.04e-6, P = 0.472; Table S6). Analyses on the subset of infected voles were not performed due to the low number of infected individuals in this dataset.

**Table 3.** Results of partial redundancy analysis (RDA) performed on the second set of bank vole samples collected at three sites: Brok, Grobka, and Długosiodło. Bank vole CFH alleles were used as explanatory variables; sampling year, site, host body mass and sex were included as covariates; and the presence or absence of *B. afzelii* OspE variants was used as response variables.

| dataset | prop.<br>conditional<br>variance | prop.<br>constrained<br>variance | adj. R<sup>2</sup> | Df | F | P value |
| --- | --- | --- | --- | --- | --- | --- |
| all<br>individuals<br>(n = 411) | 0.179 | 0.011 | -4.09E-04 | 5 | 0.968 | 0.492 |

The third dataset, containing samples collected from 12 populations across Poland, included 433 bank vole samples, of which 71 (16.40 %) were infected with *B. afzelii* (Table S2). In this dataset we identified 61 CFH/FH-R variants, including eight CFH alleles, and 33 OspE variants. The haplotype network based on OspE sequences showed that individual OspE variants were broadly shared among populations, with no clear population-specific clustering, which indicates a lack of genetic structure in the OspE gene of *B. afzelii* across the sampled populations (Figure 2). Indeed, the results of predictive CoCA showed that *B. afzelii* OspE composition does not predict vole CFH composition (Table 4). Cross-validatory % fit was negative for all CoCA axes, indicating no predictive relationship between datasets.

**Fig. 2.**
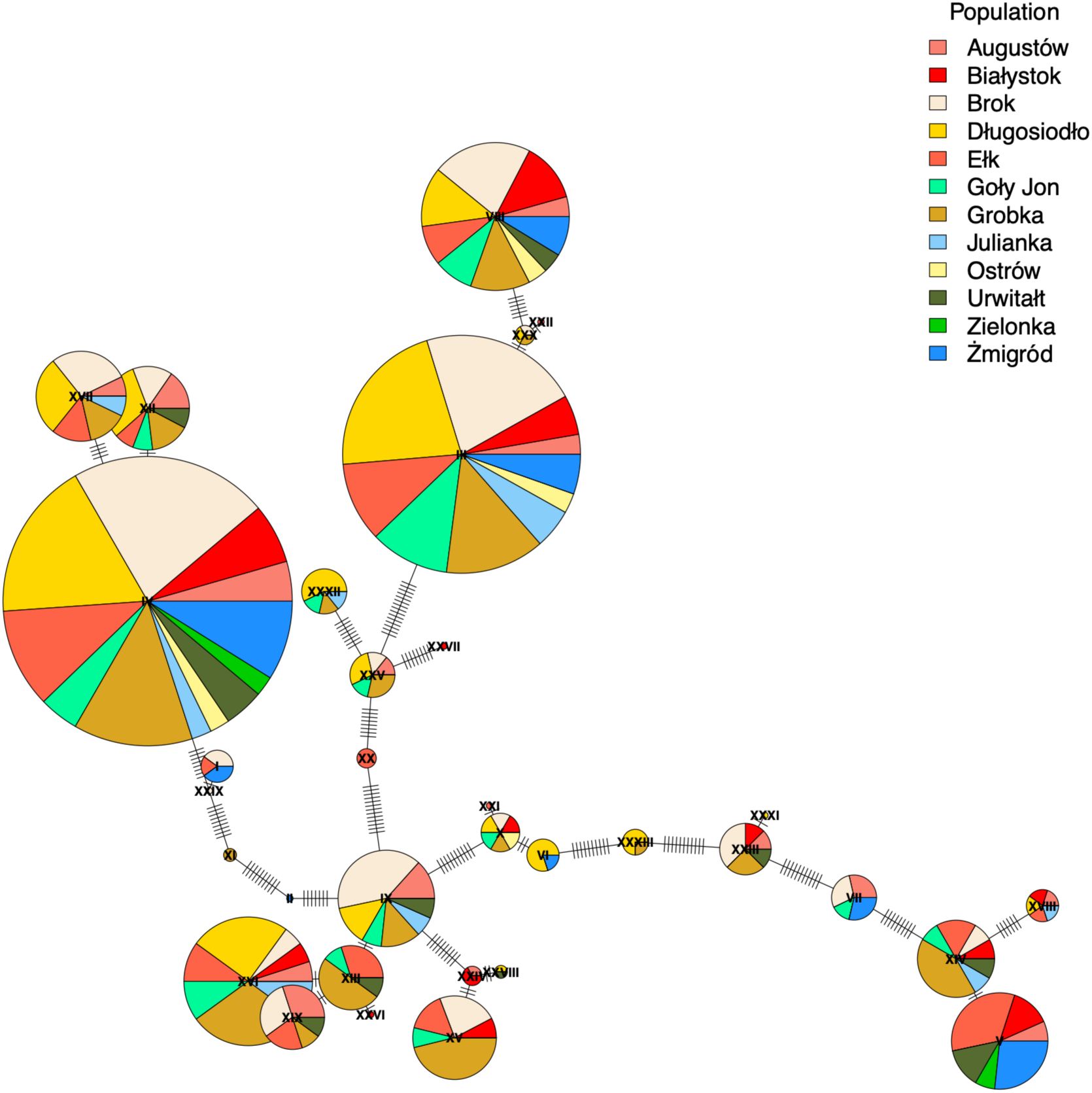
Haplotype network of *B. afzelii* OspE gene variants identified in 12 populations across Poland. Node sizes are proportional to haplotype frequencies, and pie chart colors represent the proportional contribution of each population.

**Table 4.** Results of predictive co-correspondence analysis (CoCA) performed on samples collected from 12 populations across Poland. *B.afzelii* OspE gene variant abundances in each population were used as the predictor matrix and bank vole CFH gene variant abundances in each population as the response matrix.

| axis | variance explained<br>in response (CFH) | variance explained<br>in predictor (OspE) | cross-validatory<br>%fit | P-value |
| --- | --- | --- | --- | --- |
| Comp 1 | 21.550 | 30.591 | -8.018 | 0.463 |
| Comp 2 | 21.051 | 13.982 | -20.960 | 0.664 |
| Comp 3 | 18.916 | 8.368 | -35.175 | 0.298 |
| Comp 4 | 9.587 | 8.970 | -54.728 | 0.894 |
| Comp 5 | 6.103 | 9.222 | -65.324 | 0.994 |
| Comp 6 | 7.258 | 5.729 | -203.123 | 0.884 |
| Comp 7 | 3.781 | 7.345 | -536.565 | 0.954 |

Permutation tests found that none of the axes were significant, further supporting the absence of a meaningful association between CFH and OspE. Similar results were obtained when all CFH and FH-R were included in the analysis, confirming that OspE does not predict overall bank vole CFH gene family composition (Table S7).

## Discussion

In the present study, we tested whether bank vole CFH variants are associated with infection by *B. afzelii* strains carrying different OspE variants, and whether host and pathogen genotypes exhibit spatial co-structure across populations in Poland. We found evidence that OspE evolves under positive selection at amino acids known to interact with CFH, but we did not detect significant associations between CFH genotypes and infection with specific *B. afzelii* OspE genotypes. Furthermore, we found no evidence that OspE genetic structure aligns with the structure previously reported for CFH [23].

Our study provides the first comprehensive investigation of OspE diversity and molecular evolution in *B. afzelii*. We detected 44 unique OspE variants in a total of 199 *B. afzelii*-positive samples. The variants clustered on a phylogenetic tree with *B. burgdorferi* Erp genes known to belong to the OspE subfamily (ErpA, ErpC, and ErpP; [45,46]). The tree also included two distinct clades corresponding to the OspF (ErpG, ErpK, ErpL, ErpY) and Elp (ErpB, ErpD, ErpM, ErpQ) subfamilies [46], but none of our OspE sequences clustered with those loci. Because the OspE subfamily is the only Erp group implicated in hijacking CFH [47,48], the extensive diversity we detected could possibly be a result of coevolution with the CFH of hosts supporting *Borrelia* in the wild, including bank voles.

Similarly to *B. burgdorferi* s.s., which is known to carry up to nine different Erps per strain [49], *B. afzelii* in our dataset harbored between one and six distinct OspE variants. One factor that contributes to this variation in *Borrelia* is horizontal transfer of plasmids [13,49]. We found that on the same OspC background, which is thought to nearly uniquely characterize *Borrelia* strains [50], different combinations of OspE variants occur, suggesting that horizontal OspE transfers are also common in *B. afzelii*. While this data must be interpreted with caution, as it is possible that in some cases we failed to detect coinfection, it seems unlikely that such occasional events could explain multiple cases similar to that of OspC-3a, which, in addition to OspE-20, carried unique combinations of other OspEs that were occasionally shared with different OspC variants. Horizontal transfer of OspEs appears to be the most likely explanation for these complex OspE sharing patterns (Table S4).

The diversity of OspE may be increased by frequent recombination events within the Erp gene family [13], which can be facilitated by the presence of multiple copies of OspE variants in bacterial cells, as well as those acquired by horizontal gene transfer. Indeed, omagaMap indicated evidence of recombination across the analyzed OspE sites, which may have contributed to the observed sequence variability (Table 1).

The diversity created by mutation, horizontal gene transfer and recombination may help *Borrelia* to adapt and overcome host immune defenses. In line with this, omegaMap analysis revealed evidence of positive selection acting on the ospE gene, consistent with its proposed role in immune evasion [31]. We identified eight positively selected sites (codons 116, 119, 120, 121, 122, 138, 139, and 145; Table 1). Our results overlap with findings by Cagliani et al. [24], who reported positive selection at positions 120 and 121 of the OspE protein, based on a joint analysis of several *Borrelia* species (*B. burgdorferi, B. afzelii, B. garinii, B. andersonii, B. japonica, B. lusitaniae* and *B. valaisiana*). The positively selected sites we identified are located at the known CFH-binding interface of OspE, as for example in position 120 which is known to form a hydrogen bond with CFH [18]. This supports the hypothesis that this region is subject to immune-driven selection and may play a key role in complement evasion.

However, we found no evidence of genetic associations between OspE variants and bank vole CFH alleles. Redundancy analysis, testing whether host genotype could predict the presence or absence of OspE variants, revealed no significant associations in either the full dataset or the subset of infected voles. These results suggest that OspE diversity is unlikely to be shaped solely by bank vole CFH, but it should be noted that our ability to detect genetic associations may have been limited by the relatively low infection rate in our dataset, with only 158 infected bank voles (13.79%) in the first dataset and 41 infected voles (9.98%) in the second dataset. Furthermore, the power of our analyses could have been compromised by the possible inclusion of individuals that had not been exposed to *Borrelia*. However, we note that an earlier analysis of the first sample set [28] allowed detection of even relatively small effect sizes (down to adjusted R^2^ = 1.3%) as significant. Furthermore, Różańska-Wróbel et al. [28] found that antibodies against *B. afzelii* were present in 71.10% of the individuals examined, indicating that exposure rates were high. In any case, the lack of exposure cannot explain the negative results of analyses of the infected vole subset. In this context, the very small effect sizes of CFH (adjusted R^2^ < 0.3%; Tables 2 and 3) suggest that it does not affect the probability of infection by *Borrelia* carrying specific OspE variants.

Further evidence against a tight gene × gene interaction between OspE and vole CFH variants comes from our analysis of the 12 populations sampled across Poland. For this dataset, a previous study by Notarnicola et al. [23] identified pronounced genetic structure in the bank vole CFH gene, with a clear separation between Northeastern and Southwestern Poland, a pattern we hypothesized might reflect local adaptation to *B. afzelii* strains carrying different sets of OspE genes. Such local adaptation could have evolved either within glacial refugia, or during post-glacial expansion, and could act against the flow of CFH genes between the populations, which was suggested by higher genetic differentiation between populations for CFH compared to genomic background. However, a haplotype network based on OspE sequences revealed no clear population structuring of OspE, with the vast majority of OspE variants segregating across most populations examined. Similarly, co-correspondence analysis did not reveal spatial co-structure between bank vole and *B. afzelii* genotypes. Together with the lack of individual-level associations, these spatial results further suggest that the bank vole CFH variation does not appear to be shaped by OspE composition in local *Borrelia* strains.

In the context of these negative results it is important to note that a recent study demonstrated that a *B. burgdorferi* strain lacking all cp32 plasmids, which encode Erp proteins, retains full infectivity in mice [46], indicating that OspE-mediated immune evasion may not be essential for infection in the murine host (although it may increase infection or survival success in natural situations). This also suggests that the loss of Erp proteins may be functionally compensated by other CRASPs, with overlapping roles in complement evasion, such as CspA [10]. Nevertheless, the strong signatures of positive selection we found in OspE codons interacting with CFH indicate that it is important for infection success. One possible explanation for the lack of significant association with CFH variants may be that OspE is evolving under additional selective pressures outside the system we studied, such as interactions with other hosts. While bank voles are an important reservoir for *B. afzelii*, other species such as yellow-necked mice (*Apodemus flavicollis*) and common shrews (*Sorex araneus*) may also exert selective pressure on OspE variation [25,26]. Supporting this idea, a study by Stevenson et al. [15] demonstrated that Erp proteins of *B. burgdorferi* bind to CFH, but with different specificities depending on the animal species, suggesting that *Borrelia* adapts its complement evasion mechanisms to different hosts. Additionally, a study by Hovis et al. [48] also reported that *B. burgdorferi* OspE proteins exhibit differential binding to CFH and other serum proteins from humans and various animal species. In the context of this possibility, it is remarkable that OspE exhibits very little inter-population structure, perhaps reflecting similar composition of host species in the different areas we studied. We do not have data to test this, but yellow-necked mice and shrews indeed co-occur with voles across deciduous forests in Poland [51]. Furthermore, because OspE is encoded on a multi-copy plasmid, it is possible that most *Borrelia* strains may possess a toolkit that enables infection of multiple host species. What causes the limited CFH gene flow between populations remains an open question. It could be adaptation to pathogens other than *Borrelia*. Alternatively, the expression of CFH on a genetic background that has evolved independently for a long time might result in hindered binding to self-cells, possibly compromising the function of the alternative complement pathway.

## Conclusions

In conclusion, our study provides a comprehensive analysis of the association between *B. afzelii* OspE and CFH of its rodent host, the bank vole. While we detected signatures of positive selection acting on OspE at codons capable of molecular interaction with CFH, we found no evidence of genetic or spatial associations with host CFH. These results suggest that selective pressures on bank vole CFH, which are responsible for its limited flow among populations, are not explained by *B. afzelii* OspE and likely involve additional factors. These factors should be investigated in future research.

## Declarations

## Acknowledgements

We thank Mateusz Konczal and Przemysław Gawrysiak for assistance with field work.

## Availability of data and materials

All data are available in the OSF repository: https://osf.io/e9dtb (DOI: 10.17605/OSF.IO/E9DTB).

## Authors’ contributions

JR and JRW designed the study. All authors contributed to sample collection. MG assigned vole age based on morphometric data. JRW, KP and JW performed molecular work. RFN created the reference database for CFH/FH-R distinction. JRW analysed the data, advised by JR. JRW wrote the manuscript with input and edits from JR. All authors revised the manuscript and approved the final version.

## Ethics approval

Trapping and sampling for the first dataset were approved by the First Warsaw Local Ethics Committee for Animal Experimentation (license numbers: 280/2003, 737/2007, 73/2010, and 578/2014). Trapping and sampling for the second and third datasets were approved by the Local Ethics Committee for Animal Experimentation in Poznań (decision no. 35/2021).

## References

1. Finlay BB, McFadden G. Anti-Immunology: Evasion of the Host Immune System by Bacterial and Viral Pathogens. Cell. 2006 Feb;124(4):767–82. doi:10.1016/j.cell.2006.01.034

2. Schmid-Hempel P. Immune defence, parasite evasion strategies and their relevance for ‘macroscopic phenomena’ such as virulence. Philos Trans R Soc Lond B Biol Sci. 2009;364(1513):85–98. doi:10.1098/rstb.2008.0157

3. Woolhouse MEJ, Webster JP, Domingo E, Charlesworth B, Levin BR. Biological and biomedical implications of the co-evolution of pathogens and their hosts. Nat Genet. 2002 Dec;32(4):569–77. doi:10.1038/ng1202-569

4. Arnott A, Barry AE, Reeder JC. Understanding the population genetics of Plasmodium vivax is essential for malaria control and elimination. Malar J. 2012;11:14. doi:10.1186/1475-2875-11-14

5. Mueller I, Galinski MR, Tsuboi T, Arevalo-Herrera M, Collins WE, King CL. Natural acquisition of immunity to Plasmodium vivax: epidemiological observations and potential targets. Adv Parasitol. 2013;81:77–131. doi:10.1016/B978-0-12-407826-0.00003-5

6. Rochman ND, Wolf YI, Faure G, Mutz P, Zhang F, Koonin EV. Ongoing global and regional adaptive evolution of SARS-CoV-2. Proc Natl Acad Sci U S A. 2021;118(29):2104241118. doi:10.1073/pnas.2104241118

7. Kraiczy P, Stevenson B. Complement regulator-acquiring surface proteins of Borrelia burgdorferi: Structure, function and regulation of gene expression. Ticks Tick-Borne Dis. 2013 Feb;4(1–2):26–34. doi:10.1016/j.ttbdis.2012.10.039

8. Bankhead T. Role of the VlsE Lipoprotein in Immune Avoidance by the Lyme Disease Spirochete Borrelia burgdorferi. Forum Immunopathol Dis Ther. 2016;7(3–4):191–204. doi:10.1615/ForumImmunDisTher.2017019625

9. Caine JA, Coburn J. Multifunctional and Redundant Roles of Borrelia burgdorferi Outer Surface Proteins in Tissue Adhesion, Colonization, and Complement Evasion. Front Immunol. 2016 Oct 21;7. doi:10.3389/fimmu.2016.00442

10. Lin YP, Frye AM, Nowak TA, Kraiczy P. New Insights Into CRASP-Mediated Complement Evasion in the Lyme Disease Enzootic Cycle. Front Cell Infect Microbiol. 2020 Jan 30;10:1. doi:10.3389/fcimb.2020.00001

11. Casjens S, van Vugt R, Tilly K, Rosa PA, Stevenson B. Homology throughout the multiple 32-kilobase circular plasmids present in Lyme disease spirochetes. J Bacteriol. 1997 Jan;179(1):217–27. doi:10.1128/jb.179.1.217-227.1997

12. Stevenson B. Borrelia burgdorferi erp (ospE-Related) Gene Sequences Remain Stable during Mammalian Infection. Infect Immun. 2002 Sep;70(9):5307–11. doi:10.1128/IAI.70.9.5307-5311.2002 PubMed PMID: 12183589; PubMed Central PMCID: PMC128278.

13. Brisson D, Zhou W, Jutras BL, Casjens S, Stevenson B. Distribution of cp32 Prophages among Lyme Disease-Causing Spirochetes and Natural Diversity of Their Lipoprotein-Encoding *erp* Loci. Appl Environ Microbiol. 2013 Jul;79(13):4115–28. doi:10.1128/AEM.00817-13

14. El-Hage N, Stevenson B. Simultaneous Coexpression of Borrelia burgdorferi Erp Proteins Occurs through a Specific, erp Locus-Directed Regulatory Mechanism. J Bacteriol. 2002;184(16):4536–43. doi:10.1128/jb.184.16.4536-4543.2002

15. Stevenson B, El-Hage N, Hines MA, Miller JC, Babb K. Differential Binding of Host Complement Inhibitor Factor H by *Borrelia burgdorferi* Erp Surface Proteins: a Possible Mechanism Underlying the Expansive Host Range of Lyme Disease Spirochetes. Infect Immun. 2002 Feb;70(2):491–7. doi:10.1128/IAI.70.2.491-497.2002

16. Strandh M, Råberg L. Within-host competition between *Borrelia afzelii ospC* strains in wild hosts as revealed by massively parallel amplicon sequencing. Philos Trans R Soc B Biol Sci. 2015 Aug 19;370(1675):20140293. doi:10.1098/rstb.2014.0293

17. Miller JC, Von Lackum K, Babb K, McAlister JD, Stevenson B. Temporal Analysis of *Borrelia burgdorferi* Erp Protein Expression throughout the Mammal-Tick InfectiousCycle. Infect Immun. 2003 Dec;71(12):6943–52. doi:10.1128/IAI.71.12.6943-6952.2003

18. Bhattacharjee A, Oeemig JS, Kolodziejczyk R, Meri T, Kajander T, Lehtinen MJ, et al. Structural Basis for Complement Evasion by Lyme Disease Pathogen Borrelia burgdorferi. J Biol Chem. 2013 Jun;288(26):18685–95. doi:10.1074/jbc.M113.459040

19. Meri T, Amdahl H, Lehtinen MJ, Hyvärinen S, McDowell JV, Bhattacharjee A, et al. Microbes Bind Complement Inhibitor Factor H via a Common Site. DeLeo FR, editor. PLoS Pathog. 2013 Apr 18;9(4):e1003308. doi:10.1371/journal.ppat.1003308

20. Meri S. Self-nonself discrimination by the complement system. FEBS Lett. 2016;590(15):2418–34. doi:10.1002/1873-3468.12284

21. Zipfel PF, Skerka C. Complement regulators and inhibitory proteins. Nat Rev Immunol. 2009 Oct;9(10):729–40. doi:10.1038/nri2620

22. Kenedy MR, Akins DR. The OspE-Related Proteins Inhibit Complement Deposition and Enhance Serum Resistance of *Borrelia burgdorferi*, the Lyme Disease Spirochete. Bliska JB, editor. Infect Immun. 2011 Apr;79(4):1451–7. doi:10.1128/IAI.01274-10

23. Notarnicola RF, Herdegen-Radwan M, Różańska-Wróbel J, Konczal M, Przesmycka K, Kotlík P, et al. A key regulator of missing-self innate immunity is polymorphic and under diversifying selection. Mol Biol Evol. 2026;msag082. doi:10.1093/molbev/msag082

24. Cagliani R, Forni D, Filippi G, Mozzi A, De Gioia L, Pontremoli C, et al. The mammalian complement system as an epitome of host-pathogen genetic conflicts. Mol Ecol. 2016 Mar;25(6):1324–39. doi:10.1111/mec.13558

25. Hanincová K, Schäfer SM, Etti S, Sewell HS, Taragelová V, Ziak D, et al. Association of *Borrelia afzelii* with rodents in Europe. Parasitology. 2003 Jan;126(1):11–20. doi:10.1017/S0031182002002548

26. Zhong X, Nouri M, Råberg L. Colonization and pathology of Borrelia afzelii in its natural hosts. Ticks Tick-Borne Dis. 2019 Jun;10(4):822–7. doi:10.1016/j.ttbdis.2019.03.017

27. Girard YA, Travinsky B, Schotthoefer A, Fedorova N, Eisen RJ, Eisen L, et al. Population structure of the lyme borreliosis spirochete Borrelia burgdorferi in the western black-legged tick (Ixodes pacificus) in Northern California. Appl Environ Microbiol. 2009;75(22):7243–52. doi:10.1128/AEM.01704-09

28. Różańska-Wróbel J, Migalska M, Urbanowicz A, Grzybek M, Rego ROM, Bajer A, et al. Interplay between vertebrate adaptive immunity and bacterial infectivity genes: Bank vole MHC versus *Borrelia afzelii* OSPC. Mol Ecol. 2024 Sep 24;e17534. doi:10.1111/mec.17534

29. Faith DR, Kinnersley M, Brooks DM, Drecktrah D, Hall LS, Luo E, et al. Characterization and genomic analysis of the Lyme disease spirochete bacteriophage φBB-1. PLoS Pathog. 2024;20(4):1012122. doi:10.1371/journal.ppat.1012122

30. Stevenson B, Miller JC. Intra- and Interbacterial Genetic Exchange of Lyme Disease Spirochete erp Genes Generates Sequence Identity Amidst Diversity. J Mol Evol. 2003 Sep 1;57(3):309–24. doi:10.1007/s00239-003-2482-x

31. Alitalo A, Meri T, Chen T, Lankinen H, Cheng ZZ, Jokiranta TS, et al. Lysine-Dependent Multipoint Binding of the Borrelia burgdorferi Virulence Factor Outer Surface Protein E to the C Terminus of Factor H1. J Immunol. 2004 May 15;172(10):6195–201. doi:10.4049/jimmunol.172.10.6195

32. Hellwage J, Meri T, Heikkilä T, Alitalo A, Panelius J, Lahdenne P, et al. The Complement Regulator Factor H Binds to the Surface Protein OspE of Borrelia burgdorferi. J Biol Chem. 2001 Mar;276(11):8427–35. doi:10.1074/jbc.M007994200

33. Różańska-Wróbel J, Konczal M, Notarnicola RF, Radwan J. Transcriptomic response to Borrelia afzelii infection in the skin of wild bank voles. Microbiol Spectr. 2026;14(3):0257425.

34. Liveris D, Varde S, Iyer R, Koenig S, Bittker S, Cooper D, et al. Genetic Diversity of *Borrelia burgdorferi* in Lyme Disease Patients as Determined by Culture versus Direct PCR with Clinical Specimens. J Clin Microbiol. 1999 Mar;37(3):565–9. doi:10.1128/JCM.37.3.565-569.1999

35. Kozich JJ, Westcott SL, Baxter NT, Highlander SK, Schloss PD. Development of a Dual-Index Sequencing Strategy and Curation Pipeline for Analyzing Amplicon Sequence Data on the MiSeq Illumina Sequencing Platform. Appl Environ Microbiol. 2013 Sep;79(17):5112–20. doi:10.1128/AEM.01043-13

36. Sebastian A, Herdegen M, Migalska M, Radwan J. AMPLISAS: A web server for multilocus genotyping using next-generation amplicon sequencing data. Mol Ecol Resour. 2016;16(2):498–510. doi:10.1111/1755-0998.12453

37. Pouw RB, Vredevoogd DW, Kuijpers TW, Wouters D. Of mice and men: The factor H protein family and complement regulation. Mol Immunol. 2015 Sep;67(1):12–20. doi:10.1016/j.molimm.2015.03.011

38. Wilson DJ, McVean G. Estimating diversifying selection and functional constraint in the presence of recombination. Genetics. 2006;172(3):1411–25. doi:10.1534/genetics.105.044917

39. Larsson A. AliView: A fast and lightweight alignment viewer and editor for large datasets. Bioinformatics. 2014;30(22):3276–8. doi:10.1093/bioinformatics/btu531

40. R Core Team. R: A Language and Environment for Statistical Computing. R Foundation for Statistical Computing, Vienna, Austria; 2021.

41. Posit team. RStudio: Integrated Development Environment for R. Posit Software, PBC, Boston, MA; 2024.

42. Oksanen J, Simpson G, Blanchet FG, Kindt R, Legendre P, Minchin P, et al. vegan community ecology package version 2.6-2 April 2022. 2022.

43. Paradis E. pegas: an R package for population genetics with an integrated–modular approach. Bioinformatics. 2010 Feb 1;26(3):419–20. doi:10.1093/bioinformatics/btp696

44. Simpson G. cocorresp: Co-correspondence analysis ordination methods. R Package Version 04-6. 2025.

45. Brissette CA, Cooley AE, Burns LH, Riley SP, Verma A, Woodman ME, et al. Lyme borreliosis spirochete Erp proteins, their known host ligands, and potential roles in mammalian infection. Int J Med Microbiol. 2008 Sep;298:257–67. doi:10.1016/j.ijmm.2007.09.004

46. Hillman C, Theriault H, Dmitriev A, Hansra S, Rosa PA, Wachter J. Borrelia burgdorferi lacking all cp32 prophage plasmids retains full infectivity in mice. EMBO Rep. 2025 Apr 23;26(8):1997–2012. doi:10.1038/s44319-025-00378-9

47. Alitalo A, Meri T, Lankinen H, Seppälä I, Lahdenne P, Hefty PS, et al. Complement Inhibitor Factor H Binding to Lyme Disease Spirochetes Is Mediated by Inducible Expression of Multiple Plasmid-Encoded Outer Surface Protein E Paralogs. J Immunol. 2002 Oct 1;169(7):3847–53. doi:10.4049/jimmunol.169.7.3847

48. Hovis KM, Tran E, Sundy CM, Buckles E, McDowell JV, Marconi RT. Selective Binding of *Borrelia burgdorferi* OspE Paralogs to Factor H and Serum Proteins from Diverse Animals: Possible Expansion of the Role of OspE in Lyme Disease Pathogenesis. Infect Immun. 2006 Mar;74(3):1967–72. doi:10.1128/IAI.74.3.1967-1972.2006

49. Stevenson B, Brissette CA. Erp and Rev Adhesins of the Lyme Disease Spirochete’s Ubiquitous cp32 Prophages Assist the Bacterium during Vertebrate Infection. Infect Immun. 2023;91(3):00250–22. doi:10.1128/iai.00250-22

50. Brisson D, Drecktrah D, Eggers CH, Samuels DS. Genetics of *Borrelia burgdorferi*. Annu Rev Genet. 2012 Dec 15;46(1):515–36. doi:10.1146/annurev-genet-011112-112140

51. Niedziałkowska M, Kończak J, Czarnomska S, Jędrzejewska B. Species diversity and abundance of small mammals in relation to forest productivity in northeast Poland. Écoscience. 2010;17(1):109–19. doi:10.2980/17-1-3310

