## Supplementary Material for "Does *Borrelia afzelii* outer surface protein E coevolve with complement factor H of its rodent host? Insights from GxG and spatial associations"

### Supplementary Figures

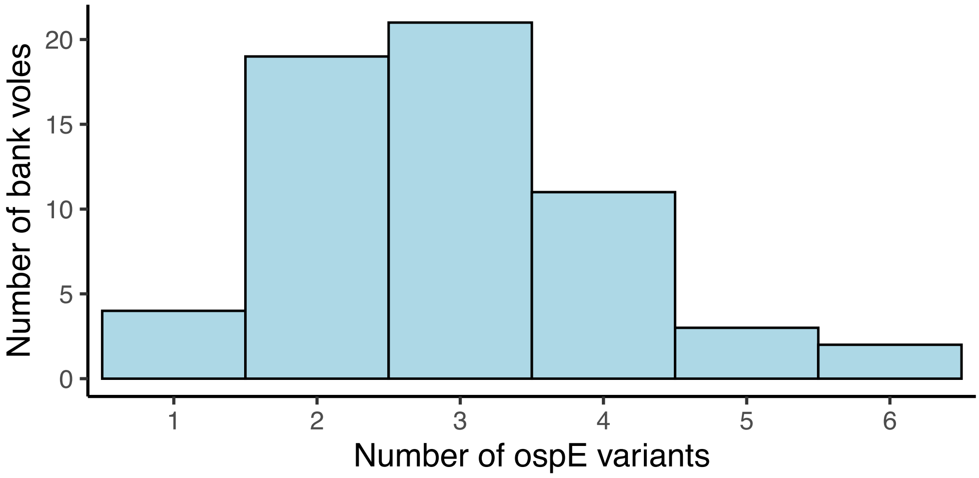


**Figure S1.** Histogram of OspE variant counts in single-strain *B. afzelii* infections identified in the dataset from Northeastern Poland.

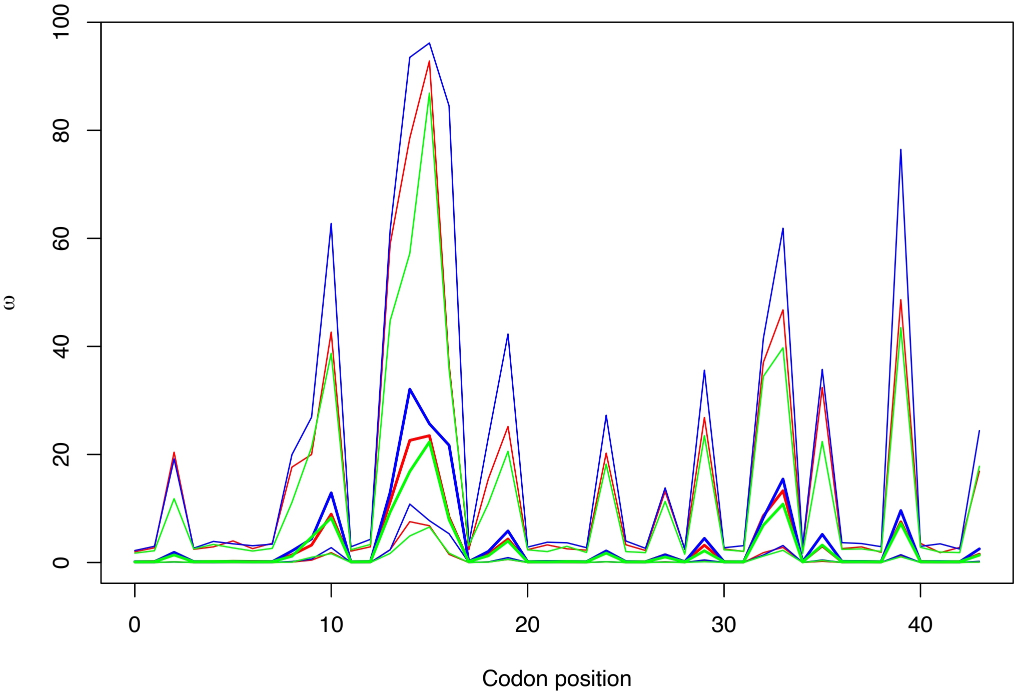


**Figure S2**. Convergence of ω (dN/dS) estimates across three independent omegaMap runs for the analyzed OspE fragment. Red, blue, and green lines correspond to runs 1, 2, and 3, respectively. Bold lines indicate posterior point estimates of ω for each run, while thinner lines represent the corresponding 95% highest posterior density (HPD) intervals.

#### Supplementary Tables

**Table S1.** First dataset used for the redundancy analysis, including populations with geographic coordinates, sampling years and sample sizes.

| Population | Coordinates | Year | # bank voles | # infected | % infected |
|---|---|---|---|---|---|
| Urwitałt | N53.48153, E21.39784 | 2002 | 71 | 7 | 9.86 |
| | | 2006 | 99 | 10 | 10.10 |
| | | 2010 | 79 | 16 | 20.25 |
| | | 2014 | 81 | 3 | 3.70 |
| | | 2018 | 59 | 4 | 6.78 |
| Tałty | N53.53644, E21.33.049 | 2002 | 68 | 5 | 7.35 |
| | | 2006 | 61 | 10 | 16.39 |
| | | 2010 | 86 | 13 | 15.12 |
| | | 2014 | 75 | 5 | 6.67 |
| | | 2018 | 69 | 5 | 7.25 |
| Pilchy | N53.42228, E21.48499 | 2002 | 72 | 18 | 25.00 |
| | | 2006 | 86 | 30 | 34.88 |
| | | 2010 | 93 | 20 | 21.51 |
| | | 2014 | 94 | 4 | 4.26 |
| | | 2018 | 53 | 8 | 15.09 |

**Table S2.** Second dataset used for the redundancy analysis, including populations with geographic coordinates, sampling years and sample sizes.

| Population | Coordinates | Year | # bank voles | # infected | % infected |
|---|---|---|---|---|---|
| Brok | N52.700283, E21.918842 | 2023 | 11 | 4 | 36.36 |
| | | 2024 | 23 | 1 | 4.35 |
| | | 2025 | 43 | 2 | 4.65 |
| Długosiodło | N52.763774, E21.632352 | 2023 | 29 | 8 | 27.59 |
| | | 2024 | 18 | 4 | 22.22 |
| | | 2025 | 52 | 2 | 3.85 |
| Grobka | N53.516669, E20.624690 | 2023 | 92 | 11 | 11.96 |
| | | 2024 | 63 | 2 | 3.17 |
| | | 2025 | 80 | 7 | 8.75 |

**Table S3.** Populations used for co-correspondence analysis and haplotype network construction, including geographic coordinates and sample sizes.

| Population | Coordinates | # bank voles | # infected bank voles |
|---|---|---|---|
| Augustów | N53.796200, E23.150766 | 34 | 5 |
| Białystok | N53.279194, E23.373034 | 23 | 4 |
| Brok | N52.700283, E21.918842 | 81 | 12 |
| Długosiodło | N52.763774, E21.632352 | 32 | 10 |
| Ełk | N53.801549, E22.399108 | 18 | 7 |
| Goły Jon | N53.693694, E18.151994 | 29 | 5 |
| Grobka | N53.516669, E20.624690 | 117 | 15 |
| Julianka | N50.773350, E19.465696 | 11 | 3 |
| Ostrów | N52.824982, E21.953556 | 3 | 1 |
| Urwitałt | N53.481530, E21.397840 | 31 | 3 |
| Zielonka | N52.582626, E17.152514 | 31 | 1 |
| Żmigród | N51.508057, E17.056494 | 23 | 5 |

**Table S4.** Haplotype combination table showing the number of OspE variant co-occurrences within samples carrying a single OspC variant, indicative of single-strain *B. afzelii* infections.

| ospC variant | ospE-1 | ospE-2 | ospE-3 | ospE-4 | ospE-5 | ospE-6 | ospE-7 | ospE-8 | ospE-9 | ospE-10 | ospE-11 | ospE-12 | ospE-13 | ospE-14 | ospE-16 | ospE-17 | ospE-18 | ospE-19 | ospE-20 | ospE-21 | ospE-23 | ospE-26 | ospE-00026 | ospE-43 | ospE-55 | ospE-00108 | Count |
|---|---|---|---|---|---|---|---|---|---|---|---|---|---|---|---|---|---|---|---|---|---|---|---|---|---|---|---|
| ospC-10b | | + | + | | | | | | + | | | | | | | | | | | + | + | | | | | | 1 |
| ospC-10b | | + | + | | | | | | | | | | | | | | | | | + | | | | | | | 1 |
| ospC-10b | | + | + | | | | | | + | | | | | | | | | | | + | | | | | | | 6 |
| ospC-10b | | + | + | | | | | | + | | | | | | | | | | | | | | | | | | 2 |
| ospC-10b | | + | + | | | | | | + | | | | | | | | | | | | | | | + | | | 1 |
| ospC-10b | + | + | | + | | | | | + | | | | | | | | | | | + | + | | | | | | 1 |
| ospC-19a | | | | | + | | | | | + | | | | + | | | | | | | | + | | | | | 1 |
| ospC-19a | | | | | + | | | | | + | | | | + | | | | | | | | | | | | | 3 |
| ospC-19a | | | | | + | | | | | + | + | | | | | | | | | | + | | | | | | 2 |
| ospC-1b | | | + | | | | | + | | | | + | | | | | | + | + | | | | | | | | 1 |
| ospC-2a | | | | | | | | | | + | | | | + | | | | | | | | + | | | | | 1 |
| ospC-2a | | | + | | | | | | | | + | | | + | | | | | | | | | | | | | 1 |
| ospC-3a | + | | | | | | | | | | | | | | | + | | | + | | | | | | | | 2 |
| ospC-3a | | | | | | | | | | | | | | | | + | | | + | | | | | | | | 1 |
| ospC-3a | | | + | | | | | | | | | | | | | | | | + | | | | | | | | 2 |
| ospC-3a | | | | | | | | | | | | | + | | | | | | + | | | | | | | | 1 |
| ospC-3a | | | | | | | | | | | | | | | | | | | + | | | | | | | + | 1 |
| ospC-3a | + | | | | | | | | | | | | + | | | | | | + | | + | | | | | | 1 |
| ospC-3a | | | + | | | | | | | | | | | | | | | | + | | + | | | | | | 1 |
| ospC-3a | | | | | | | | | | | | | | | | | | | + | | | | | | | | 2 |
| ospC-4c | | | + | | | | | + | + | | | | | | | + | | | | + | | | | | | | 1 |
| ospC-4c | | + | + | | | | | | | | | | | | | | | | | | | | | | | | 1 |
| ospC-4c | | | + | | | | | | | | | | | | | | | | | | | | | | | | 2 |
| ospC-4c | | | | | | | | | | + | | | | | + | | | | | | | | | | + | | 1 |
| ospC-4c | | | | | | | | | | + | | | | | + | | | | | | | | | | | | 1 |
| ospC-4c | | | + | | | | | + | | | | | | | | | | | | | | | | | | | 2 |
| ospC-4c | + | | | | | | + | + | + | | | | | | | | + | | | | | | + | | | | 1 |
| ospC-4c | | | + | | | | | | | | | | | | + | | | | | | | | | | | | 1 |
| ospC-4c | | | | | | | | | | | | | | | + | | | | | | | | | | + | | 1 |
| ospC-4c | | | + | | | | | | | | | + | | | | | | | | | | | | | | | 1 |
| ospC-7b | + | | | | | + | + | | | | | | | | | | | | | | | | | | | | 6 |
| ospC-7b | + | | | | | + | | | | | | | | | | | | | | | | | | | | | 4 |
| ospC-7b | | | | | | + | + | | | | | | | | | | | | | | | | | | | | 2 |
| ospC-9c | + | | | | | | | | | | | | | | | | + | | | | | | | | | | 1 |
| ospC-9c | + | | | + | | | | | | | | | | | | | + | | | | | | | | | | 3 |

**Table S5.** Results of partial redundancy analysis (RDA) performed on the first set of bank vole samples collected at three sites: Urwitałt, Tałty, and Pilchy. Bank vole CFH and FH-R variants were used as explanatory variables; sampling year, site, host age and sex were included as covariates; and the presence or absence of *B. afzelii* OspE variants was used as response variables.

| dataset | prop. conditional variance | prop. constrained variance | adj. R2 | Df | F | P value |
|---|---|---|---|---|---|---|
| all individuals (n = 1146) | 0.050 | 0.010 | 1.81E-04 | 12 | 1.018 | 0.424 |
| infected individuals (n = 158) | 0.100 | 0.062 | -0.013 | 12 | 0.835 | 0.897 |

**Table S6.** Results of partial redundancy analysis (RDA) performed on the second set of bank vole samples collected at three sites: Brok, Grobka, and Długosiodło. Bank vole CFH and FH-R variants were used as explanatory variables; sampling year, site, host body mass and sex were included as covariates; and the presence or absence of *B. afzelii* OspE variants was used as response variables.

| dataset | prop. conditional variance | prop. constrained variance | adj. R2 | Df | F | P value |
|---|---|---|---|---|---|---|
| all individuals (n = 411) | 0.179 | 0.039 | 7.04E-06 | 17 | 1.000 | 0.472 |

**Table S7.** Results of predictive co-correspondence analysis (CoCA) performed on samples collected from 12 populations across Poland. *B.afzelii* OspE variant abundances in each population were used as the predictor matrix and bank vole CFH/FH-R variant abundances in each population as the response matrix.

| axis | variance explained in response (CFH) | variance explained in predictor (OspE) | cross-validatory %fit | P-value |
|---|---|---|---|---|
| Comp 1 | 10.817 | 34.165 | -1.867 | 0.433 |
| Comp 2 | 16.374 | 13.599 | -5.478 | 0.289 |
| Comp 3 | 11.045 | 10.989 | -6.571 | 0.775 |
| Comp 4 | 11.011 | 8.505 | -7.741 | 0.849 |
| Comp 5 | 8.800 | 7.235 | -9.116 | 0.990 |
| Comp 6 | 7.474 | 7.134 | -8.680 | 0.998 |
| Comp 7 | 9.018 | 4.718 | -10.725 | 0.948 |
| Comp 8 | 5.770 | 4.734 | -11.154 | 0.991 |
| Comp 9 | 5.759 | 4.459 | -12.785 | 0.959 |
| Comp 10 | 6.578 | 3.155 | -10.354 | 0.536 |
| Comp 11 | 5.766 | 1.309 | -22.371 | 0.010 |
